# Bayesian modelling of high-throughput sequencing assays with malacoda

**DOI:** 10.1101/819466

**Authors:** Andrew R. Ghazi, Xianguo Kong, Ed S. Chen, Leonard C. Edelstein, Chad A. Shaw

**Affiliations:** Quantitative and Computational Biosciences, Baylor College of Medicine, Houston, Texas; Cardeza Foundation for Hematologic Research, Thomas Jefferson University, Philadelphia, Pennsylvania; Molecular and Human Genetics, Baylor College of Medicine, Houston, Texas

## Abstract

NGS studies have uncovered an ever-growing catalog of human variation while leaving an enormous gap between observed variation and experimental characterization of variant function. High-throughput screens powered by NGS have greatly increased the rate of variant functionalization, but the development of comprehensive statistical methods to analyze screen data has lagged behind. In the massively parallel reporter assay (MPRA), short barcodes are counted by sequencing DNA libraries transfected into cells and output RNA in order to simultaneously measure the shifts in transcription induced by thousands of genetic variants. These counts present many statistical challenges, including over-dispersion, depth dependence, and uncertain DNA concentrations. So far, the statistical methods used have been rudimentary, employing transformations on count level data and disregarding experimental and technical structure while failing to quantify uncertainty in the statistical model.

We have developed an extensive framework for the analysis of NGS functionalization screens available as an R package called malacoda (available from github.com/andrewGhazi/malacoda). Our software implements a probabilistic, fully Bayesian model of screen data. The model uses the negative binomial distribution with gamma priors to model sequencing counts while accounting for effects from input library preparation and sequencing depth. The method leverages the high-throughput nature of the assay to estimate the priors empirically. External annotations such as ENCODE data or DeepSea predictions can also be incorporated to obtain more informative priors – a transformative capability for data integration. The package also includes quality control and utility functions, including automated barcode counting and visualization methods.

To validate our method, we analyzed several datasets datasets using malacoda and alternative MPRA analysis methods. These data include experiments from the literature, simulated assays, and primary MPRA data. We also used luciferase assays to experimentally validate the strongest hits from our primary data, as well as variants for which the various methods disagree and variants detectable only with the aid of external annotations.

**Author Summary:** Genetic sequencing technology has progressed rapidly in the past two decades. Huge genomic characterization studies have resulted in a massive quantity of background information across the entire genome, including catalogs of observed human variation, gene regulation features, and computational predictions of genomic function. Meanwhile, new types of experiments use the same sequencing technology to simultaneously test the impact of thousands of mutations on gene regulation. While the design of experiments has become increasingly complex, the data analysis methods deployed have remained overly simplistic, often relying on summary measures that discard information. Here we present a statistical framework called for the analysis of massively parallel genomic experiments designed to incorporate prior information in an unbiased way. We validate our method by comparing our method to alternatives on simulated and real datasets, by using different types of assays that provide a similar type of information, and by closely inspecting an example experimental result that only our method detected. We also present the method’s accompanying software package which provides and end-to-end pipeline that provides a simple interface for data preparation, analysis, and visualization.

## Introduction

The advent of next generation sequencing (NGS) has generated an explosion of observed genetic variation in humans. Variants with unclear effects greatly outnumber those with obvious, severe impact; the 1000 Genomes Project [1] has estimated that a typical human genome has roughly 150 protein-truncating variants, 11,000 peptide-sequence altering variants, and 500,000 variants falling into known regulatory regions. Simultaneously, genome-wide association studies (GWAS) have found strong statistical associations between thousands of noncoding variants and hundreds of human phenotypes [2,3]. Traditional methods of assessing the regulatory impact of variants are slow and low-throughput: luciferase reporter assays require multiple replications of cloning individual genomic regions, transfection into cells, and measurement of output intensity.

Massively Parallel Reporter Assays (MPRA), overviewed in Figure 1, were developed to assess simultaneously the transcriptional impact of thousands of genetic variants [4]. The simplest form of MPRA uses a carefully designed set of barcoded oligonucleotides containing roughly 150 base pairs of genomic context surrounding variants of interest. There are typically thousands of variants selected by preliminary evidence from GWAS, and there are usually ten to thirty replicates of each allele with different barcodes. The oligonucleotides are cloned into plasmids, making a complex library that is then transfected into cells. The cells use the library as genetic material and actively transcribe the inserts. Because the barcodes are preserved by transcription, counting the RNA products of each variant construct by re-identifying each barcode in the NGS product provides a direct measure of the transcriptional output of a given genetic variant. By designing the oligonucleotide library to contain multiple barcodes of both the reference and alternate alleles for each variant, one can statistically assess the transcription shift (TS) for each variant.

**Fig 1.**
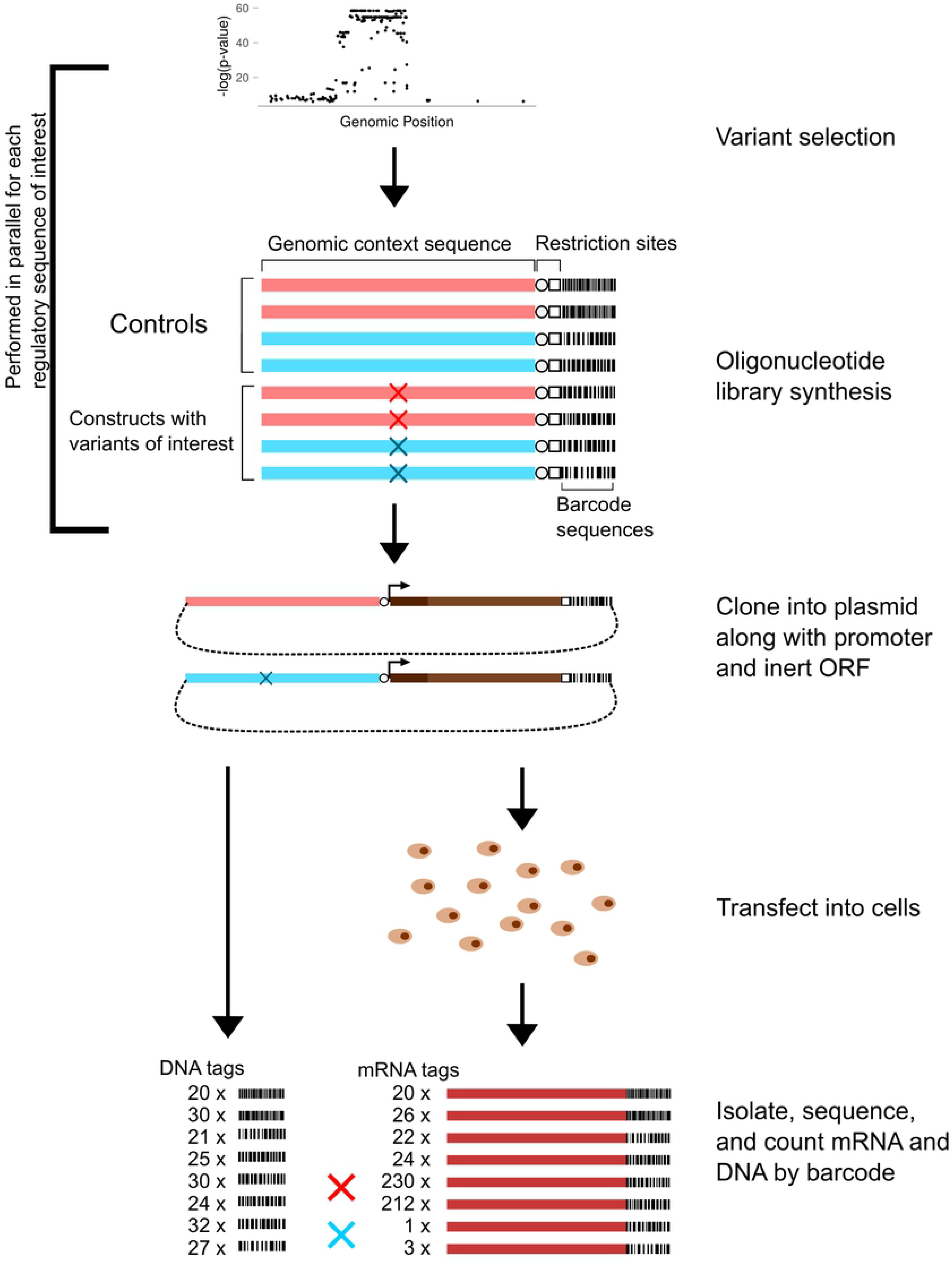
Diagram of MPRA. MPRA simultaneously assess the transcription shift of thousands of variants. The diagram shows six constructs with two variants, but in practice the size of the oligonucleotide library is only limited by cost. A typical MPRA has tens to hundreds of thousands of oligonucleotides to assay thousands of variants.

MPRA have successfully identified many transcriptionally functional variants [5, 6, 7], but the accompanying statistical analyses have been rudimentary. Initial studies focused on the computation of the “activity” for each barcode in each RNA sample. This involves averaging across depth-adjusted counts to compute a normalizing DNA factor for each barcode, then dividing RNA counts by the DNA factor and taking the log of this ratio. Then a t-test is used to compare the activity measurements for each allele, followed by assay-wide multiple-testing corrections. The key limitations include ignoring systematic variation due to unknown DNA concentrations, the application of heavy transformation and summarization to the data prior to modelling, and the failure to include the reservoir of prior data and biological knowledge concerning genes and genomic regions. The methods mpralm [8] and MPRAscore [9] are more recent methods, but they suffer from a number of limitations: failure to model variation in input DNA concentrations, aggregation of data across barcodes and sequencing samples without modeling systematic sources of variation, and over-reliance on point estimates of dispersion that cause systematic errors in transcription shift estimates.

Other areas of genomic analysis have generated a wealth of information on genomic structure and function, frequently specific to particular genomic contexts and variants. For example, the ENCODE project [10] provides genome-wide ChIP-seq data on transcription binding profiles, histone marks, and DNA accessibility. Computational methods such as DeepSea [11] use machine learning to provide variant-specific predictions on chromatin effects. Genome-wide databases like ENCODE and computational predictors like DeepSea contain real information about variant effects, but the method for incorporating this information into a statistical framework for experimental analysis of variants has been unclear.

We hypothesized that a Bayesian approach to high throughput NGS screens such as MPRA would improve statistical sensitivity and specificity and yield more accurate estimates of variant function, particularly when incorporating prior information. The Bayesian approach offers a flexible modeling system that can flexibly fit hierarchical model structures of count data while also directly accounting for experimental sources of variation. The Bayesian approach also enables the integration of prior information and probabilistic modelling of dispersion parameters. These advantages offer significant improvements in statistical efficiency and provide advantages for formulating systems-level hypotheses -- for example, the impact of specific transcription factors -- that are absent from other approaches. Here we present *malacoda*, an end-to-end Bayesian statistical framework that addresses the gaps in the prior approaches while providing novel methods for incorporating prior information. The malacoda method centers on MPRA but also has potential extension to a broad array of NGS-based high-throughput screens. We establish the superior performance of malacoda on MPRA compared to alternatives using simulation studies. Then, we apply the method to previously published findings to make new biological discoveries that we explore in the paper. We also apply malacoda to primary MPRA studies that we performed. The results demonstrate that using malacoda we can discover biologically important findings that were missed by prior approaches. We have made the software available as an open source R package on GitHub.

## Methods

### Overview

In malacoda we utilize a negative binomial model for NGS to consider barcode counts with empirically estimated gamma priors, and we explicitly model variation in the input DNA concentrations for each barcode. By default the method marginally estimates the priors from the maximum likelihood estimates of each variant in the assay; the method also supports informative prior estimation by using external genomic annotations for each variant as weights. This approach enables disparate knowledge sources to inform the results in a principled, systematic, and calculation. The probabilistic model underlying malacoda uses the NGS data directly without transformation, and it accounts for all known sources of experimental variation and uncertainty in model parameters. Finally, the method provides estimate shrinkage as a method for avoiding false positives.

### Description of the statistical model

MPRA data are the counts of the barcoded DNA input from sequencing the plasmid library and counts of the barcoded RNA outputs from sequencing the RNA content extracted from passaged cells. The DNA counts vary according to the sequencing depth of the sample as well as due to the inherent noise in library preparation. The RNA measurements also vary according to sequencing depth, but they are also affected by the DNA input concentration and the inherent transcription rate of their associated region of genomic context. Figure 2A shows a subset of a typical MPRA dataset, with two barcodes of each allele for two variants and several columns of counts. We find that typically MPRA are performed with four to six RNA sequencing replicates and a smaller number of DNA replicate samples. Figure 2B shows a simplified Kruschke diagram of the model underlying malacoda, using the mean-dispersion parameterization of the negative binomial. More explicitly,

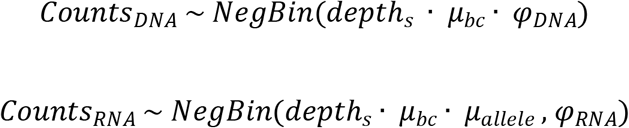

Where depth_s_ indicates the depth of a particular sequencing sample, μ_bc_ indicates the unknown concentration of a particular barcode in the plasmid library, and μ_allele_ indicates the effect of the genomic context of a given allele of a given variant. There are separate dispersions parameters φ for both DNA and RNA. The means μ and dispersions φ come from their own gamma priors.

**Fig 2.**
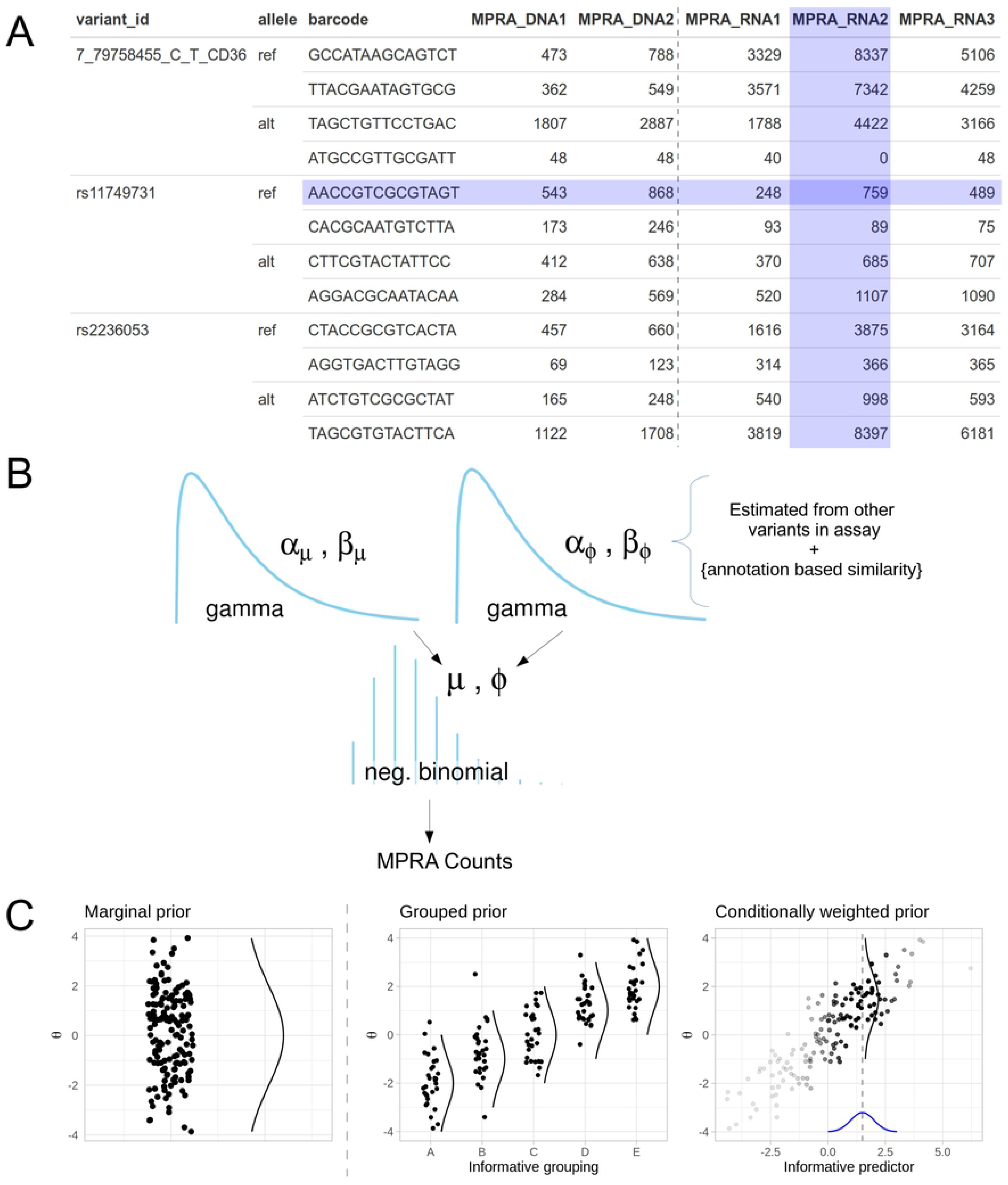
MPRA data and malacoda priors. A) The table shows a subset of our primary MPRA data. Highlighted cell containing 759 is influenced both by the sequencing depth of its sample (column) and the unknown input DNA concentration of its barcode (row). B) A simplified Kruschke diagram of the generative model underlying malacoda C) A conceptual diagram demonstrating three prior types available from malacoda. The marginal prior (left) weights all variants in the assay equally, while the grouped and conditional priors utilize informative annotations as weights in the prior estimation process.

The negative binomial distribution is a natural choice for modelling NGS count data given its ability to accurately fit overdispersed observations frequently seen in sequencing data [12]. Briefly, the observed dispersion in NGS count data usually exceeds that expected from simpler binomial or Poisson models. We chose gamma distributions as priors for several reasons. They have the appropriate [0,∞) support, and for a non-negative random variable whose expectation and expected log exist, they are the maximum entropy distribution. Additionally, they are characterized by two parameters, allowing the prior estimation process to accurately fit the observed population of negative binomial estimates. Probabilistic modelling of the dispersion parameters is key -- as demonstrated by simulation in S1 Appendix. This practice helps avoid pitfalls common to methods based on point estimates of dispersion parameters. The barcode-level count data model is a central contribution of the malacoda method.

After computing the joint posterior on all model parameters, the posterior on transcription shift is computed as a generated quantity by taking the difference between log means of the alternate and reference alleles. 95% highest density interval on TS is used to make binary calls on whether a variant is functional or non-functional. If the interval excludes zero as a credible value, the variant is labelled as functional. An optional “region of practical equivalence” can be defined on a per-assay basis when there is particular interest in rejecting transcription shift values around zero [13].

### Empirical priors

The gamma priors are fit empirically by maximum likelihood estimation. Specifically, each variant-level model is first fit by maximizing the likelihood component of the malacoda model, then gamma distributions are fit to those estimates for the means and dispersions of the DNA, reference RNA, and alternate RNA. This approach offers several benefits. First, it leverages the high-throughput nature of the assay. The full dataset determines the prior; in situations with thousands of variants the individual contribution of each variant to the prior is negligible. Secondly, it constrains the prior to be reasonable in the context of a given assay. Specific circumstances regarding library preparation, sequencer properties, cell culture conditions, and other unknown factors will cause the underlying statistical properties of each MPRA to be unique. A less informed, general-purpose prior, such as gamma(α = .001, β = .001), would assign a considerable amount of probability density to unreasonable regions of parameter space. Empirical estimation ensures that the priors capture the reasonable range of values for each parameter while avoiding putting unwarranted density on extreme values [14]. Finally, by sharing information between variants, empirical priors provide estimate shrinkage. The prior effectively regularizes all parameter estimates, a behavior which is important in multi-parameter models with relatively little data per parameter. This in turn acts as a natural safeguard against false positives, thus removing the need for *post hoc* multiple testing correction.

In order to incorporate external knowledge, the malacoda method also allows users to provide arbitrary annotations to supplement the analysis. Figure 2C contrasts the marginal prior estimation (left) with two prior types that make use of external annotations. These priors make use of the information in the annotations by employing the principle that similarly annotated variants should perform similarly in the assay. When the annotations are simply a set of descriptive categories (for example predictions of likely benign, uncertain, or likely functional), the grouped prior (2C, center) simply fits a prior distribution within each subset. When the annotations are continuous values, the conditionally weighted (2C, right) prior employs a kernel smoothing process to estimate the prior. To estimate the prior for a single variant, it initializes a t-distribution kernel centered at the annotation of the variant in question, then gradually widens this kernel until the *n*-th most highly weighted variant (where *n* is a configurable tuning parameter defaulting to 100) has a weight of at least one percent of that of the most influential variant. While the diagram in figure 2C shows this for only a single informative annotation on the horizontal axis, the code allows for an arbitrary number of continuous predictors to be used.

### Simulation and Validation Studies

We took several approaches to validate and compare the malacoda method with alternatives. First, we simulated MPRA data using across a realistic grid of parameters governing the fraction of truly functional variants, the number of variants in the assay, and the number of barcodes per allele. These simulations also modelled distinct sequencing samples, varying sequencing depth, and barcode failure during library preparation. We then compared malacoda to alternative methods including the t-test, mpralm, and MPRAscore. Across these simulations we compared performance metrics such as area-under-curve (AUC) and estimate accuracy. Secondly, we applied malacoda and alternative methods to real MPRA data from the Ulirsch dataset [5], using inter-method consensus as a performance metric. We repeated this with our own primary MPRA data on variants related to platelet function. Finally, we tested a subset of variants where the various methods disagreed with luciferase reporter assays to assess consistency with MPRA estimates of variant function.

### Software

Our method is available as an R package from github.com/andrewGhazi/malacoda. The package includes detailed installation instructions, extensive help documentation, an analysis walkthrough vignette, and implementations of traditional activity-based analysis methods. The package also includes functionality to extract, quality-filter, and count barcodes from a set of FASTQ files through an application of the FASTX-Toolkit [15]. Through an interface with the FreeBarcodes package [16], the package can also decode sequencer errors in the barcodes of an assay that has been designed using our previous work, mpradesigntools [17]. In our experience this typically recaptures about 5% additional data with no additional cost beyond a line of code during the assay design. The package also contains plotting functionality to help visualize the results of analyses.

## Experimental Methods

In order to collect experimental measurements of the transcriptional impact of variants through means other than MPRA, we performed luciferase reporter assays on sixteen variants. Four were among the strongest signals detected in our MPRA, six were variants from our MPRA where the statistical methods disagreed, and six were variants from the Ulirsch dataset [5] where the malacoda marginal and DeepSea-based [11] conditional prior model fits disagreed.

150-200bp genomic DNA sequences flanking the variants were amplified by PCR using K562 lymphoblast (ATCC) genomic DNA as template, then cloned into PGL4.28 minimum promoter luciferase reporter vector (Promega) at NheI and HindIII sites. Counterpart SNP variants were generated by site-directed mutagenesis. All the constructs were validated by DNA sequencing. 3µg plasmid preparations were co-transfected with 0.5μg β-gal plasmid into 1×10^6^ of K562 cells with Lipofectamine 2000 based on manufacturer's instructions. Each assay was repeated with 3 independent plasmid preparations. 24 hours post transfection, luciferase and β-gal were measured. Luciferase units were then normalized to β-gal values.

## Results

### Simulation Studies

We evaluated our simulation results in three ways. First, we focused on the accuracy of transcription shift estimates. Figure 3A shows the results of analyzing one simulated dataset, with the true value of the simulation’s transcription shift plotted on the x-axis, with the model estimates on the y-axis. For each fit of each simulation using each analysis method, we analyzed performance using two metrics: standard deviation of estimates for truly non-functional variants at zero (center dots, lower is better) and correlation with the truth for truly functional variants with nonzero effects (off-center dots, higher is better).

**Fig 3.**
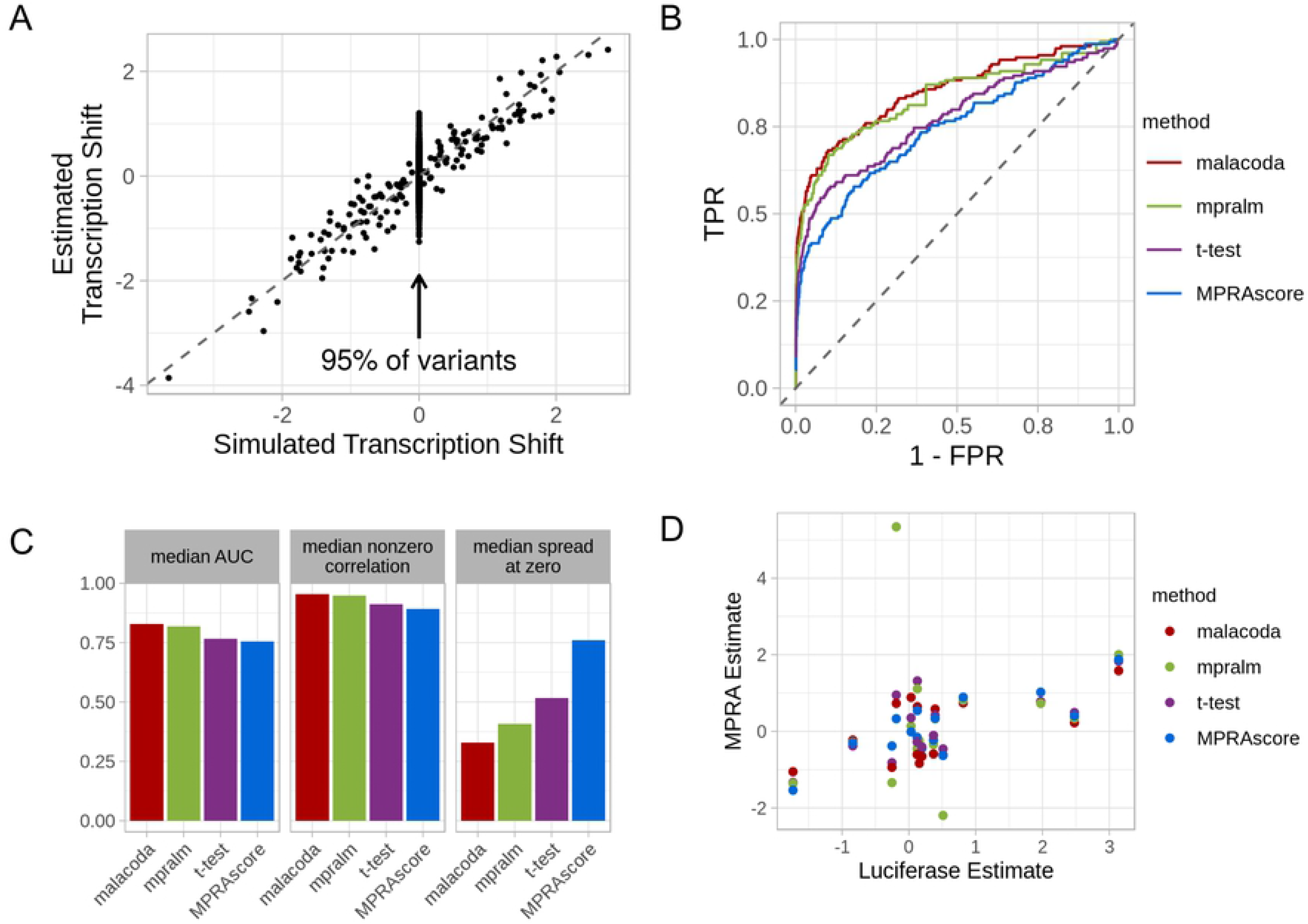
Simulation results. A) The figure compares TS values used to generate simulated data to malacoda TS estimates. Simulated MPRA assays use a varying fraction of variants that are truly non-functional (center). B) ROC curves assess the performance of each method on a randomly selected assay with 3000 variants, 5% truly functional variants, and 10 barcodes per allele. C) Performance metrics averaged across multiple simulations under the same conditions as B. D) A scatterplot demonstrates the relationship between luciferase-based estimates of TS against MPRA-based estimates.

Second, we also computed area under the curve (AUC) for each method. Bayesian methods such as malacoda explicitly do not consider a null hypothesis and therefore do not output p-values; in order to create an analogous output quantity to derive an ROC curve we instead computed one minus the minimum HDI width necessary to include zero as a credible transcription shift value to distinguish true and false positives. Figure 3B shows the ROC curves by method for a randomly chosen simulation with ten barcodes per allele, 5% truly functional variants, and 3000 variants. Figure 3C shows that across all simulations with these characteristics, malacoda consistently showed the highest median AUC, highest correlation with the truth for functional variants, and the lowest spread among estimates of truly nonfunctional variants. Other simulation grid points are shown in S2 Appendix, and these display similar patterns.

In order to examine the performance of malacoda on real data, we applied the various methods to both the Ulirsch data [5] and to our own primary dataset. Unlike the case with simulations, the underlying true values are not known. However, inter-method consensus can serve as a performance metric -- alternative methods presumably fail in different ways, so if they tend to disagree with one another but agree with malacoda, that would imply that malacoda is working well across the cases where others fail. Indeed, Figure 4 shows that the other methods tend to correlate with malacoda better than the other alternatives. The one exception is when applied to our dataset, mpralm tends to agree best with the t-test method. Given that linear models underlie both mpralm and the t-test method, it seems plausible that they would sometimes show similar results.

**Fig 4.**
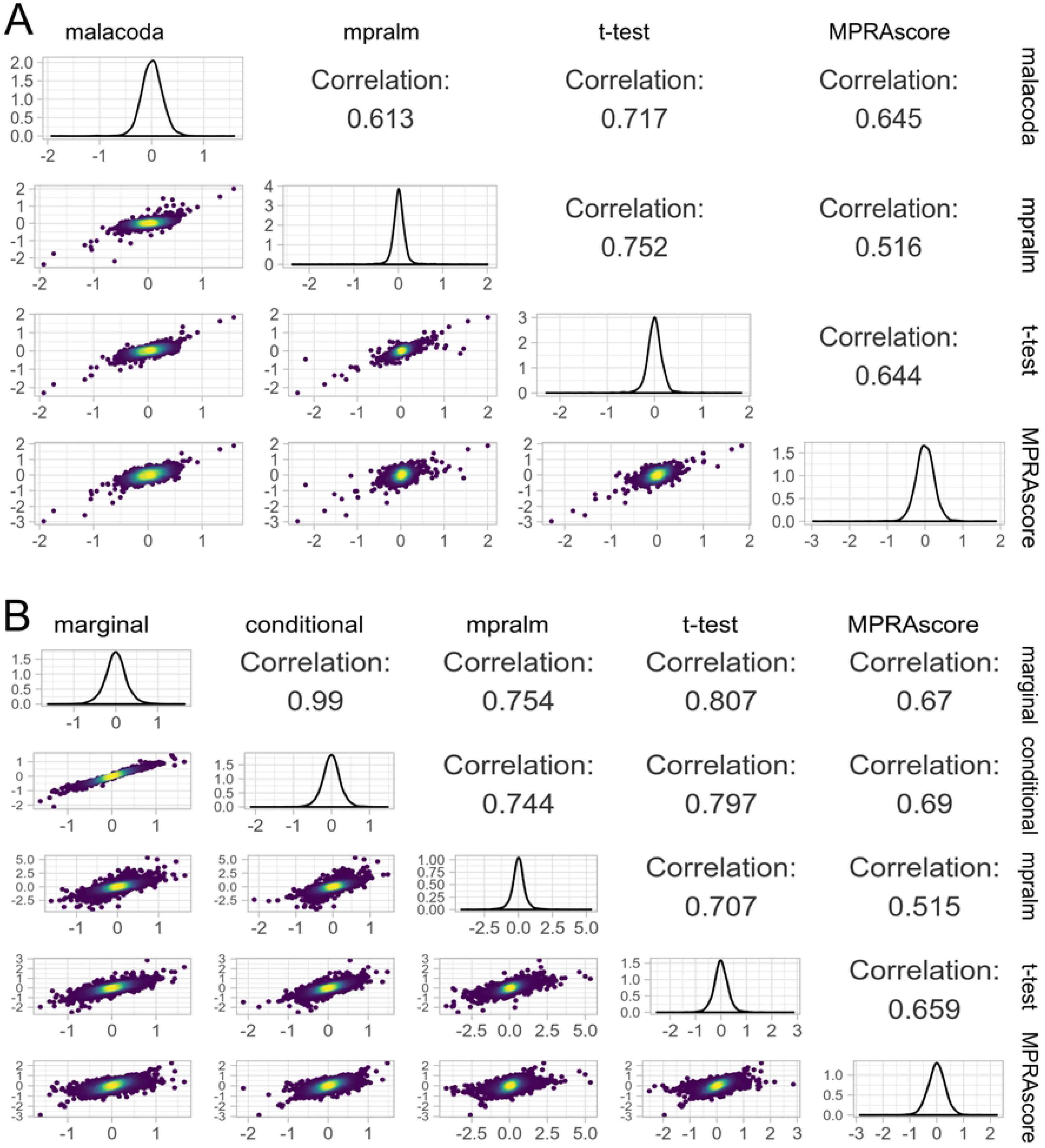
Inter-method consensus. A) A pairwise plot of TS estimates in our MPRA, showing that other methods generally agree with malacoda more than each other. Color indicates local density of points. B) A pairwise plot of TS estimates using both the marginal and DeepSea-based malacoda priors in the Ulirsch dataset, showing a similar outcome.

### Biological results

The number of luciferase reporter assays we performed was not enough to overcome the amount of noise inherent to light intensity-based measurements, thus we did not have enough data to clearly demonstrate that any of the MPRA analysis methods outperform the others in terms of correlation with luciferase results. However, the results show that the various methods are consistent with MPRA-based estimates Figure 3D, providing further evidence that MPRA results are biologically realistic.

We closely inspected a particular biological discovery to demonstrate malacoda’s ability to identify low-signal variants. One of the functional variants we identified with malacoda using the DeepSea-based conditional prior in the Ulirsch dataset [5] is rs11865131; this variant is identified by malacoda but not by any of the other methods. The variant rs11865131 is in an intron within the *NPRL3* gene which encodes the Natriuretic Peptide Receptor Like 3 protein. *NPRL3* is part of the GTP-ase activating protein activity toward Rags [18] (GATOR1) complex. The GATOR1 complex inhibits mammalian target of rapamycin (*MTOR*) by inhibiting *RRAGA* function (reviewed in [18] *MTOR* signaling has been implicated in platelet aggregation and spreading in addition to aging associated venous thrombosis [19, 20]. Analysis of the rs11865131 locus indicates that it colocalizes with ENCODE ChIP-Seq peaks for 36 transcription factors in K562 erytholeukemia cells as well as containing enhancer histone epigenetic marks. Together, these data indicate that this is likely an important regulatory region. In addition to the heterologous K562 cell line, data from cultured megakaryocytes indicates that rs11865131 lies within *RUNX1* and *SCL* ChIP-Seq peaks, two well-studied megakaryopoietic transcription factors [21]. This agrees with our data that platelet *NPRL3* mRNA is positively associated with platelet count in healthy humans [22, 23]. These data indicate that malacoda has identified a likely important regulatory region for megakaryocytes and platelets that was missed by other MPRA analysis methods.

## Discussion

We developed a fully Bayesian framework for the analysis of NGS high throughput screens with specific application to MPRA studies. The method is an advance in statistical and computational science for these data - a fully Bayesian model that probabilistically incorporates all known sources of variation. The method does a better job of identifying true positives in simulated data and performs well in empirical studies. The method identified a previously missed functional variant in the *NPRL3* gene that has confirmatory evidence from a variety of other studies. Particular advantages of the method are accurate estimation of variant effects, the treatment of the dispersion parameter in both estimation and inference, and the potential to incorporate informative prior information.

The functional discovery of the variant rs11865131 represents a demonstration of the power of the malacoda method to identify biologically important results missed by alternative methods. This variant lies in an intronic region of the gene *NPRL3*, and protein coding approaches to variant analysis would overlook this regulatory variant. Multiple lines of evidence point to the biological relevance of this variant, including epigenetic and transcription factor binding data as well as evidence of association with platelet count in healthy humans.

There are downsides to our method. First, Bayesian methods that estimate a joint posterior on many parameters by MCMC are significantly slower than optimization approaches. To address this, we fit our models with Stan [24], which allows us to perform a first pass fit with Automatic Differentiation Variational Inference [25] and, if seemingly worthwhile, to perform a final fit with Stan’s state-of-the-art No-U-Turn Sampler. Despite this measure, our marginal prior analysis of 8251 variants from the Ulirsch dataset with 50,000 MCMC samples using no variational first pass took over fifteen hours when parallelized across eighteen threads on two Intel Xeon X5675 3.07GHz processors. Nevertheless, an analysis that runs in hours is reasonable for an assay that takes weeks to perform.

Secondly, the efficacy of our method does not account for uncertainty in our empirical prior estimation functionality [14]. The R package includes a fully hierarchical model that adds an additional layer of hyperparameters in order to probabilistically model the gamma priors and all other parameters for an entire MPRA dataset at once, but this approach falls outside the intended scope of the malacoda framework. This model, featuring hundreds of thousands of parameters, is presently too complex to fit in practice.

The statistical method and validation work presented in this article has focused primarily on the analysis of “typical” MPRA: two alleles per variant, in a single tissue type, with no other experimental perturbations. However, we have expanded the modelling capabilities of the package beyond these limitations. Models tailored to more exotic experimental structures, such as arbitrary numbers of alleles per variant, multiple tissue types, or cell-culture perturbations, are also included with the package. We also have expanded the model framework included in the package beyond MPRA into CRISPR screen modelling: the counts of gRNAs targeting specific genes in survival/dropout screens can make use of an analogous negative binomial structure with similar empirical gamma priors. This opens the path to incorporating gene-level annotations into Bayesian CRISPR screen analysis.

Sophisticated high-throughput assays are a central component to the future of genomics. Therefore, the statistical methods used for these data should be as efficient as possible, accounting for all sources of variation and quantifying the resulting uncertainty. Our software, malacoda, provides an end-to-end framework for the probabilistic analysis of MPRA data. Through our well-documented, easy-to-use R package, users can perform sequencing error correction and data pre-processing before executing a fully Bayesian analysis in as little as two lines of code. When informative annotations on variant function are available, malacoda is capable of taking full advantage through a conditional prior estimation process. We hope that this work may act as a stepping stone towards further integrative, probabilistic analysis in the field of high-throughput genomics.

## Supporting Information

**S1 Appendix. Negative Binomial variance estimation.**

**S2 Appendix. Simulation details and extended results.**

**S3 Dataset. RData file of luciferase and MPRA results**. An RData file that loads two objects: luc_results, a table of the luciferase results, and mpra_results, giving the primary data on MPRA counts for the variants tested with luciferase

**S4 Dataset. RData file of estimate comparisons.**The data necessary to produce Figure 4. An RData file that contains two data frames: ulirsch_comparisons and primary_comparisons. Each row corresponds to one variant, and each column corresponds to a given analysis method. The values in the table give the transcription shift estimates.

## References

1. Auton A, Abecasis GR, Altshuler DM, Durbin RM, Bentley DR, Chakravarti A, et al. A global reference for human genetic variation. Nature [Internet]. 2015;526(7571):68–74 doi: 10.1038/nature15393

2. Hindorff LA, Sethupathy P, Junkins HA, Ramos EM, Mehta JP, Collins FS, et al. Potential etiologic and functional implications of genome-wide association loci for human diseases and traits. Proc Natl Acad Sci USA. 2009;106(23):9362–7 doi: 10.1073/pnas.0903103106

3. Nishizaki SS, Boyle AP. Mining the Unknown: Assigning Function to Noncoding Single Nucleotide Polymorphisms. Trends Genet [Internet]. 2017;33(1):34–45 doi: 10.1016/j.tig.2016.10.008

4. Melnikov A, Murugan A, Zhang X, Tesileanu T, Wang L, Rogov P, et al. Systematic dissection and optimization of inducible enhancers in human cells using a massively parallel reporter assay. Nat Biotechnol [Internet]. 2012;30(3):271–7 doi: 10.1038/nbt.2137

5. Ulirsch JC, Nandakumar SK, Wang L, Giani FC, Zhang X, Rogov P, et al. Systematic functional dissection of common genetic variation affecting red blood cell traits. Cell [Internet]. 2016;165(6):1530–45 doi: 10.1016/j.cell.2016.04.048

6. Tewhey R, Kotliar D, Park DS, Liu B, Winnicki S, Reilly SK, et al. Direct identification of hundreds of expression-modulating variants using a multiplexed reporter assay. Cell [Internet]. 2016;165(6):1519–29 doi: 10.1016/j.cell.2016.04.027

7. Shen SQ, Myers CA, Hughes AEO, Byrne LC, Flannery JG, Corbo JC. Massively parallel cis-regulatory analysis in the mammalian central nervous system. Genome Res. 2016;26(2):238–55 doi: 10.1101/gr.193789.115

8. Myint L, Avramopoulos DG, Goff LA, Hansen KD. Linear models enable powerful differential activity analysis in massively parallel reporter assays. BMC Genomics. 2019;20(1):1–19 doi: 10.1186/s12864-019-5556-x

9. Niroula A, Ajore R, Nilsson B. MPRAscore: robust and non-parametric analysis of massively parallel reporter assays. Bioinformatics. 2019;(July):1–3 doi: 10.1093/bioinformatics/btz591

10. Consortium EP. An integrated encyclopedia of DNA elements in the human genome. Nature. 2013;489(7414):57–74 doi: 10.1038/nature11247.An

11. Zhou J, Troyanskaya OG. Predicting effects of noncoding variants with deep learning-based sequence model. (DeepSea). Nat Methods [Internet]. 2015;12(10):931–4 doi: 10.1038/nmeth.3547

12. Love MI, Huber W, Anders S. Moderated estimation of fold change and dispersion for RNA-seq data with DESeq2. Genome Biol [Internet]. 2014;15(12):550. doi: 10.1186/s13059-014-0550-8

13. Kruschke J. Doing Bayesian Data Analysis: A Tutorial with R, JAGS, and Stan. 2nd ed. London: Academic Press; 2015. P.336–40.

14. Gelman A, Carlin JB, Stern HS, Dunson DB, Vehtari A, Rubin DB. Bayesian Data Analysis. Third Edition. Boca Raton, FL: CRC Press; 2013. p. 51-6, p. 102–4.

15. Assaf G, Hannon GJ. FASTX-Toolkit [Internet]. 2010. Available from: http://hannonlab.cshl.edu/fastx_toolkit/index.html

16. Hawkins JA, Jones SK, Finkelstein IJ, Press WH. Indel-correcting DNA barcodes for high-throughput sequencing. Proc Natl Acad Sci [Internet]. 2018;115(27):E6217–26 doi: 10.1073/pnas.1802640115

17. Ghazi AR, Chen ES, Henke DM, Madan N, Edelstein LC, Shaw CA. Design tools for MPRA experiments. Bioinformatics. 2018;34(15):2682–3 doi: 10.1093/bioinformatics/bty150

18. Shaw RJ. GATORs take a bite out of mTOR. Science. 2013;340(6136):1056–7 doi: 10.1126/science.1240315

19. Aslan JE, Tormoen GW, Loren CP, Pang J, McCarty OJT. S6K1 and mTOR regulate Rac1-driven platelet activation and aggregation. Blood. 2011;118(11):3129–36 doi: 10.1182/blood-2011-02-331579

20. Yang J, Zhou X, Fan X, Xiao M, Yang D, Liang B, et al. MTORC1 promotes aging-related venous thrombosis in mice via elevation of platelet volume and activation. Blood. 2016;128(5):615–24 doi: 10.1182/blood-2015-10-672964

21. Chacon D, Beck D, Perera D, Wong JWH, Pimanda JE. BloodChIP: A database of comparative genome-wide transcription factor binding profiles in human blood cells. Nucleic Acids Res. 2014;42(D1):172–7 doi: 10.1093/nar/gkt1036

22. Simon LM, Edelstein LC, Nagalla S, Woodley AB, Chen ES, Kong X, et al. Human platelet microRNA-mRNA networks associated with age and gender revealed by integrated plateletomics. Blood. 2014;123(16):37–45 doi: 10.1182/blood-2013-12-544692

23. Edelstein LC, Simon LM, Montoya RT, Holinstat M, Chen ES, Bergeron A, et al. Racial differences in human platelet PAR4 reactivity reflect expression of PCTP and miR-376c. Nat Med [Internet]. 2013;19(12):1609–16 doi: 10.1038/nm.3385

24. Carpenter B, Gelman A, Hoffman MD, Lee D, Goodrich B, Betancourt M, Brubaker M, Guo J, Li P, Riddell A. Stan: A probabilistic programming language. J Stat Softw. 2017;76(1):. doi: 10.18637/jss.v076.i01

25. Kucukelbir A, Blei DM, Gelman A, Ranganath R, Tran D. Automatic Differentiation Variational Inference. J Mach Learn Res. 2017;18:1–45 Available from: https://arxiv.org/abs/1603.00788

